# Development of spontaneous firing of fusiform neurons from the dorsal cochlear nucleus of mice occurs after hearing onset

**DOI:** 10.1101/2021.09.06.459172

**Authors:** Nikollas M. Benites, Beatriz Rodrigues, Carlos H. Silveira, Ricardo M. Leão

## Abstract

The dorsal cochlear nucleus (DCN) in the auditory brainstem integrates auditory and somatosensory information. Mature fusiform neurons express two qualitative intrinsic states in equal proportions: quiet, with no spontaneous regular action potential firing, or active, with regular spontaneous action potential firing. However, how these firing states and other electrophysiological properties of fusiform neurons develop during early postnatal days to adulthood is not known. Thus, we recorded fusiform neurons from mice from P4 to P21 and analyzed their electrophysiological properties. In the pre-hearing phase (P4-P13), we found that fusiform neurons are mostly quiet, with the active state emerging after hearing onset at P14. Subthreshold properties present more variations before hearing onset, while action potential properties vary more after P14, developing bigger, shorter, and faster action potentials. Interestingly, the activity threshold is more depolarized in pre-hearing cells suggesting that persistent sodium current (I_NaP_) increases its expression after hearing. In fact, I_NaP_ increases its expression after hearing, accordingly with the development of active neurons. Thus, we suggest that the post-hearing expression of I_NaP_ creates the active state of the fusiform neuron. At the same time, other changes refine the passive membrane properties and increase the speed of action potential firing of fusiform neurons.

## Introduction

The electrophysiological activity of mature neurons in the central nervous system is different from that displayed during development. The biophysical properties of the ionic currents of neurons undergo constant change during the period of neuronal development. Consequently, the behavior of individual neurons and the networks in which they are part is different from behavior seen in the adult state. The biophysical properties of these currents, such as voltage dependence, kinetics, and the subunits that make up the ion channel, can change during development (Moody & Bosma, 2005). However, the complex mechanisms by which the different biophysical properties of subliminal currents establish the different neuronal phenotypes are not fully understood. For example, several neurons fire spontaneously during the embryonic and postnatal period, and this spontaneous activity disappears in adulthood (Moody & Bosma, 2005). Transient expression of ionic currents during development that disappears later in adulthood is also common, e.g., sodium and potassium currents disappear in adult retinal and cochlear neurons (Zhou & Fain, 1996); Marcotti et al., 1999).

The rodent auditory brainstem neurons undergo several physiological and morphological changes in the postnatal period culminating around the 12^th^-14^th^ postnatal day when the ear canal opens, and the animal begins to hear. Several experiments in the Medial Nucleus of the Trapezoid Body (MNTB) showed that this region undergoes several alterations during the pre-hearing period. For example, pre-synaptic action potentials become shorter during development, as do pre-synaptic sodium currents (Taschenberger & Von Gersdorff, 2000; Leão *et al*., 2005*b*), and the calyceal terminals progressively increase their ability to trigger action potentials at high frequency without failing, a consequence of a faster recovery from the inactivation of these channels (Taschenberger & Von Gersdorff, 2000; Leão *et al*., 2005*b*). The synaptic also vesicles become more coupled to calcium channels, increasing their likelihood of exocytosis (Taschenberger *et al*., 2002; Leão & von Gersdorff, 2009). These alterations, among others, occur before the opening of the auditory canal at P12, suggesting that they are pre-programmed alterations and not triggered by sensory experience. However, sensory hearing deprivation is known to cause subtle but essential changes in excitability and neurotransmission in the MNTB and bushy cells in the anteroventral cochlear nucleus (Leao *et al*., 2004, 2006; Leão *et al*., 2005*b*; Grande *et al*., 2014; Clarkson *et al*., 2016; Zhuang *et al*., 2017). Thus, it seems that, at least in these synapses, most of the alterations originate from genetic programming. However, sensory-dependent post-hearing changes occur that are relevant for the fine-tuning of the auditory processing.

The dorsal cochlear nucleus (DCN) is part of the cochlear nuclei that are the first central station of the auditory pathway and integrates acoustic information with multimodal sensory signals that come from the most diverse areas of the brain (Fujino & Oertel, 2003; Oertel & Young, 2004; Pilati *et al*., 2012*a*) and its neural pathways assume to detect spectral cues for localizing sounds (May, 2000; Young & Davis, 2002; Oertel & Young, 2004) as well to filter out self-generated noise (Shore & Zhou, 2006; Singla *et al*., 2017). The fusiform (or principal) neuron of the DCN integrates the afferent synaptic inputs to the DCN, projecting axons to the inferior colliculus (Cant & Benson, 2003). The fusiform neurons exhibit a regular firing pattern when stimulated with a constant depolarizing current (Pilati *et al*., 2012*b*). Furthermore, numerous *in vitro* and *in vivo* studies have shown that fusiform neurons exist in one of two qualitatively different intrinsic states: a quiet state, where the membrane potential remains close to a stable resting membrane potential (RMP), or a spontaneously active state, where action potentials are spontaneously fired (Leao *et al*., 2012). Close to 50% of fusiform present spontaneous (active neurons) firing, while the other 50% do not fire spontaneous (quiet neurons) (Pilati *et al*., 2012*b*; Leao *et al*., 2012; Zugaib *et al*., 2016).

The active firing state is created by the expression of a persistent sodium current (I_NaP_) that lowers the threshold for action potential firing and an inward-rectifying potassium current that sets the membrane potential above or below the threshold set for I_NaP_ (Leao *et al*., 2012; Ceballos *et al*., 2016). These two states increase the dynamic range of fusiform neurons allowing these neurons to decrease firing in response to acoustic stimulation (Young & Davis, 2002). However, the development of this heterogeneity is unknown. Here we aim to study the development of the intrinsic electrophysiological properties of fusiform neurons of the DCN from the pre-hearing period to the post-hearing period. We found that the active state develops after hearing onset in parallel with the expression of I_NaP_. We hypothesized that the expression of I_NaP_ is regulated by hearing and creates the firing diversity seen in these neurons

## Materials and Methods

### Animals

We used Swiss mice of both sexes from postnatal days 4 to 21 (P4-21). On most occasions, pregnant females were monitored each day to check the day of birth of the pups. Animals were kept in a 12/12 hours dark/light cycle with food and water *ad libtum*. The ethics committee animal experimentation of the University of São Paulo approved all procedures (CEUA. Protocol 133/2018).

### Auditory brainstem response (ABR)

For recording ABRs, we used an RZ6 multi-processor (Tucker Davis Technology, USA) connected to a speaker (Multi-Field Magnetic Speaker, Tucker Davis Technology), which were previously calibrated in the open field configuration. Animals were anesthetized with ketamine-xylazine (60mg/kg and 10mg/kg, respectively, i.p.) after a sedative dose of isoflurane. The animals were laid on a non-electrical heating pad inside an electrically and acoustically isolated chamber. Three platinum needle electrodes (impedance ∼1 kΩ) were placed in the head of the animals in the following way: a reference electrode, bellow the pina ipsilateral to the stimulus; a ground electrode, placed at the contralateral pina; and a recording electrode at the head vertex. Electrodes were connected to a RA4PA medusa preamplifier (Tucker Davis Technology, USA) and connected to the RZ6 multi-processor system via an optic fiber. Data were acquired with the BioSigRZ software (25 kHz of the sampling rate). A 0.1 ms, single-channel mono-phasic click protocol was presented at a rate of 21/s from 90 to 20 dB SQL in 10 dB steps. The software averaged 512 neural responses to each stimulus presentation with a gain set at 20 dB. Data were filtered from 0.3 to 3 kHz and stored for offline analysis. The threshold was considered as the lowest intensity level where sound stimulus-evoked any wave peak could be recognized.

### Brainstem slice preparation

Mice were anesthetized with isoflurane and decapitated. The brainstems were being removed after cutting the vestibulocochlear nerve. The brainstems were dipped in melted agarose (2,5%) at 38 ° C, and the temperature was quickly reduced with a metal block to solidify the agarose. The piece was mounted in a compresstome (VF300-0Z, Precisionary Instruments, USA), and coronal brain slices containing the dorsal cochlear nucleus, DCN, (200 µm thick) were cut, in an ice-cold solution containing (in mM) 87 NaCl, 2.5 KCl, 1.25 NaH_2_PO_4_, 25 NaHCO_3_, 0.2 CaCl_2_, 7 MgCl_2_, 25 dextrose, and 75 sucrose, pH 7.4 when oxygenated with 95% O_2_/5% CO_2_. After, the slices were incubated in artificial cerebrospinal fluid (aCSF) for 35 minutes at 35°C and then at room temperature (25°C) for electrophysiological experiments. The aCSF containing (in mM) 120 NaCl, 2.5 KCl, 1.25 NaH_2_PO_4_, 25 NaHCO_3_, 2 CaCl_2_, 1 MgCl_2_, and 10 dextrose, pH 7.4 when oxygenated with 95% O_2_/5% CO_2_ (osmolality 310 mOsmol/kg H_2_O).

### Electrophysiology

The DCN and fusiform cells were localized under oblique illumination with a microscope (BX51W, Olympus, Japan). Recordings were obtained with borosilicate microelectrodes (BF150-86-10, Sutter Instrument Company, USA) with resistance between 2 and 4 MΩ when filled with internal solution (in mM) 130 K-gluconate, 20 KCl, 10 Na_2_-phosphocreatine, 10 HEPES, 0.1 EGTA, 2 Mg-ATP, and 0.2 Na-GTP, pH 7.4 adjusted with KOH, with a final osmolality of 295 mOsmol/kg H_2_O. Fusiform cells were identified by their location at the principal layer in the DCN and biophysical characteristics (Tzonopoulos et al., 2003). Whole-cell patch-clamp recordings were performed using an EPC-10 patch-clamp amplifier (Heka Elektronics, Germany) under continuously aCSF perfusion (∼1mL/min) and temperature (31-34°C) controlled using an in-line heater (Warner Instruments). Whole-cell recordings started after at least 5 minutes after attaining whole-cell configurations. Series resistance was monitored constantly during the recordings, and neurons with >20 MΩ were discarded. Series resistance compensation was compensated between 50 and 80%

Recordings were performed in the presence of strychnine (2 µM), picrotoxin (100 µM), and DNQX (10 µM). Quiet neurons were neurons with no action potential firing at rest, or sparse, the unregular firing of less than 1 Hz (Leão et al., 2011). Resting membrane potential was measured in the presence of tetrodotoxin (TTX; Alomone Labs, Israel) (0.5 µM). Voltage-current relationships were performed by injecting 1s current pulses with 50 pA steps from -200 pA. In active neurons, DC negative current (−20 pA to -200 pA) was injected to stop the action potentials firing.

Voltage-clamp recordings were performed at a holding potential of -65 mV with voltage membrane from -105 mV to -55mV in 10mV steps. Data were acquired using the PATCHMASTER (Heka) at a 50 kHz rate and filtered at a low-pass filter (3 kHz, Bessel).

### Data analysis

All recordings were corrected offline by subtracting a measured liquid junction potential of -10mV. Data were analyzed using routines written in Igor Pro (Wavemetrics, Portland, USA). We measure the RMP using the modal voltage determined by an all-points histogram from a stretch of 52 seconds. Input resistance was measured by analyzing the slope of the VI relationship at steady-state (last 100 ms) and at the peak of the hyperpolarization. The membrane time constant was obtained fitting a single exponential function on the hyperpolarizing decay during a -50 pA pulse. The depolarization sag was calculated as the difference between the peak and the steady-state hyperpolarization produced by a -200 pA current. Rheobase is defined as minimal current necessary for firing an action potential from the rest or in active neurons from a potential close to -70 mV. The activity threshold was calculated as in Leão et al. (2011) as the mean membrane potential at the current step right before the start of spontaneous action potential firing.

Action potential parameters were measured using a phase plane plot (dV/dt vs. V) from the first action potential at rheobase. The threshold was considered the potential in the phase plane, at which the slope reached 10 V.s^-1^. Action potential amplitude was calculated as the difference between the threshold and the action potential peak calculated in the phase plane plot as the more depolarized point of 0 V/s. The maximum rate of rise (ROR) was calculated as the peak value in the phase plane plot.

Half-width was the action potential duration at half of the amplitude, and the latency was the time between the start of the stimulus and the threshold of the first action potential at rheobase. Fast afterhyperpolarization (fAHP) was the difference between the threshold and the more hyperpolarized potential after action potential repolarization.

Data are presented as the mean ± SEM. We compared the differences in means using Student t-tests, one-way analysis of variance (ANOVA) with a Fischer’s post-test, correlation, and linear regression performed in GraphPad Prism 9.0 (GraphPad Software, USA) with a significance level set at p<0.05. The principal component analysis was performed using a routine written in R.

## Results

### The proportion of active neurons increases after hearing onset

Rodents start to hear after 9 to 14 days after birth (Alford & Ruben, 1963; Hack, 1968; Ehret, 1976; Sonntag *et al*., 2009). In order to establish the precise hearing onset of our mice, we examined mice from P12 to P14 in litters we knew the exact day of birth of the pups. We observed that before P14, mice did not respond with a startle in response to a clap or whistle. Additionally, we detected the opening of the ear canal at P14. ABR recordings in response to clicks (Figure 1A) showed that mice at P12 did not respond to clicks up to 90 dB, while at P13, we observed some response of the wave I near 50 dB (50 ± 18.37 dB; n=4). At P14 the waves were more evident and at lower thresholds (Figure 1A), which decreased even further at P18 and P21 (P14: 28.7 ± 1.1 dB; P18: 23.2 ±1,2 dB; P21 23.8 ± 1 dB; n = 19, 19 and 13 respectively. p = 0.001, one-way ANOVA; Figure 1B). We then assumed a hearing onset at P14 in our animals.

**Figure 1.**
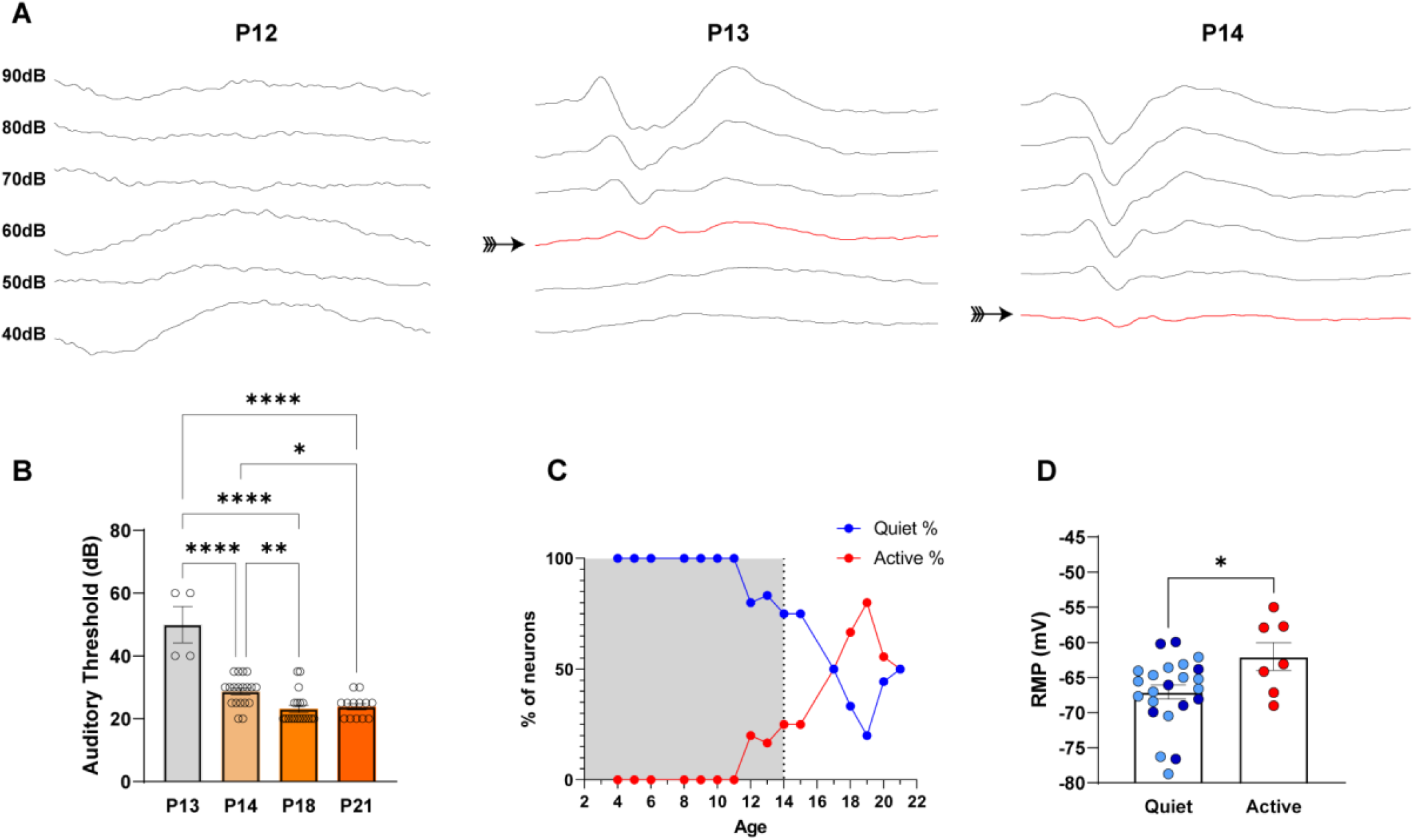
Determination of hearing onset. **A**. Auditory brainstem responses of a P12, P13, and P14 mice. The arrow shows the hearing threshold. **B**. Hearing thresholds at different ages **C**. Proportion between quiet and active neurons across ages. The grey box represents the pre-hearing phase **D**. RMP in the presence of TTX light blue circles are data from rpe-hearing neurons and dark circles from post-hearing neurons. p <0.05 *, p<0.01 **, p<0.001 ***, p<0.0001 ****.

We performed whole-cell patch-clamp recordings in fusiform neurons from mice across 4 to 21 postnatal days (P4-P21). Neurons from 4-13 days old animals were classified as pre-hearing and animals P14 and above as post-hearing. We found that quiet fusiform neurons were predominant in pre-hearing animals (95.92%, 47 out of 49 cells), and the proportion of active and quiet fusiform neurons is similar after hearing onset (quiet: 51.52%, 16 out of 33 neurons; active 48.48%, 17 out of 33 neurons) as previously reported (Leao *et al*., 2012; Zugaib *et al*., 2016). We found that fusiform neurons were all quiet until P11, with some active neurons detected at P12 and P13. After P14, the proportion of active neurons gradually increased, reaching a peak at P19 (Figure 1C). We conclude that the active state of the fusiform neuron is characteristic of the post-hearing phase.

A fundamental difference of active and quiet neurons was a more depolarized resting membrane potential (RMP) of active neurons, which were the main factor responsible for the existence of the two firing modes of fusiform neurons (Leao *et al*., 2012). We confirmed this observation in our mice when we measured the RMP of active and quiet neurons in the presence of TTX, being the active neurons more depolarized than the quiet neurons (quiet: -67.1 ± 1 mV; active: -62.0 ± 2 mV, n = 23 and 7 respectively; p = 0.02 unpaired t-test. Figure 1D). The RMP of quiet neurons was not different from pre or post-hearing mice (pre-hearing: -67.2 ± 1.2 mV; post-hearing: -66.7 ± 1.9 mV; n = 15 and 8 respectively; p = 0.8 unpaired t-test). We confirmed then that active fusiform neurons from our mice have more depolarized RMP than quiet neurons.

### Pre-hearing quiet neurons are less excitable than post-hearing neurons

We then compared the action potential firing of pre-hearing and post-hearing neurons. Pre-hearing neurons fire fewer action potentials in response to depolarization currents than active or post-hearing quiet neurons (Figure 2Bi). In response to a 250 pA current pre-hearing quiet neurons fired 41 ± 10 action potentials, while post-hearing active neurons fired 71 ± 26 action potentials and active neurons fired 62 ± 15 action potentials (one-way ANOVA: p=0.010, pre-vs. post-quiet: p=0.008; pre-quiet vs. active: p=0.04; post-quiet vs.active: p=0.48; Table 1).

**Table 1.** ANOVA post-tests data statistics. The n values for variables were 47 of pre-quiet, 17 of post-quiet, and 18 of active. Some n values were different: resting membrane potential, n = 15 of pre-quiet, 8 of post-quiet, and 7 of active; the number of action potentials at 250 of pre-quiet, post-quiet, and active were 37,11 and 14, respectively; current at -55 mV, n = 39, 15, and 16 for pre-, post-quiet, and active, respectively; I_NaP_ n= 14, 5, and 7 fot pre-, post-quiet, and active, respectively. The bold values highlight α<-0.05.

|  | Uncorrected Fisher's LSD | Mean 1 | Mean 2 | Mean diff. | p-value | 95.00% CI of diff. |
| --- | --- | --- | --- | --- | --- | --- |
| Activity threshold (mV) | Pre-quiet vs. post-quiet | -56.96 | -62.25 | 5.297 | **0.0009** | 2.248 to 8.347 |
|  | Pre-quiet vs. active | -56.96 | -66.20 (18) | 9.242 | **<0.0001** | 6.256 to 12.23 |
|  | Post-quiet vs. active | -62.25 | -66.20 | 3.945 | **0.0342** | 0.3013 to 7.589 |
| AP Amplitude ($\Delta$ mV) | Pre-quiet vs. post-quiet | 76.47 | 90.63 | -14.16 | **0.0001** | -21.13 to -7.192 |
|  | Pre-quiet vs. active | 76.47 | 96.94 | -20.47 | **<0.0001** | -27.29 to -13.64 |
|  | Post-quiet vs. active | 90.63 | 96.94 | -6.304 | 0.1359 | -14.63 to 2.025 |
| AP Latency (s) | Pre-quiet vs. post-quiet | 0.2180 | 0.05552 | 0.1624 | **0.0073** | 0.04498 to 0.2799 |
|  | Pre-quiet vs. active | 0.2180 | 0.04341 | 0.1745 | **0.0034** | 0.05951 to 0.2896 |
|  | Post-quiet vs. active | 0.05552 | 0.04341 | 0.01211 | 0.8641 | -0.1282 to 0.1525 |
| AP Threshold (mV) | Pre-quiet vs. post-quiet | -42.65 | -43.88 | 1.229 | 0.5298 | -1.491 to 3.949 |
|  | Pre-quiet vs. active | -42.65 | -45.76 | 3.110 | **0.0180** | 0.4459 to 5.774 |
|  | Post-quiet vs. active | -43.88 | -45.76 | 1.881 | 0.3550 | -1.369 to 5.132 |
| fAHP ($\Delta$ mV) | Pre-quiet vs. post-quiet | 19.62 | 25.00 | -5.378 | **0.0022** | -8.768 to -1.989 |
|  | Pre-quiet vs. active | 19.62 | 26.88 | -7.256 | **<0.0001** | -10.58 to -3.937 |
|  | Post-quiet vs. active | 25.00 | 26.88 | -1.878 | 0.3588 | -5.928 to 2.172 |
| Current at -55mV | Pre-quiet vs. post-quiet | 106.3 | 39.99 | 66.34 | 0.0281 | 7.361 to 125.3 |
|  | Pre-quiet vs. active | 106.3 | -85.35 | 191.7 | <0.0001 | 134.0 to 249.3 |
|  | Post-quiet vs. active | 39.99 | -85.35 | 125.3 | 0.0006 | 55.57 to 195.1 |
| Halfwidth (ms) | Pre-quiet vs. post-quiet | 0.4249 | 0.2192 | 0.2057 | **0.0002** | 0.1013 to 0.3101 |
|  | Pre-quiet vs. active | 0.4249 | 0.1973 | 0.2275 | **<0.0001** | 0.1253 to 0.3298 |
|  | Post-quiet vs. active | 0.2192 | 0.1973 | 0.02184 | 0.7284 | -0.1029 to 0.1466 |
| $I_{NaP}$ (-55mV) | Pre-quiet vs. post-quiet | -26.16 | -86.86 | 60.70 | 0.0941 | -11,21 to 132,6 |
|  | Pre-quiet vs. active | -26.16 | -161.4 | 135.3 | **0.0002** | 71,38 to 199,2 |
|  | Post-quiet vs. active | -86.86 | -161.4 | 74.57 | 0.0688 | -6,244 to 155,4 |
| Max. Rate of rise ($V.s^{-1}$) | Pre-quiet vs. post-quiet | 570.3 | 952.8 | -382.5 | **<0.0001** | -519.0 to -246.1 |
|  | Pre-quiet vs. active | 570.3 | 1036 | -465.3 | **<0.0001** | -599.0 to -331.6 |
|  | Post-quiet vs. active | 952.8 | 1036 | -82.75 | 0.3156 | -245.8 to 80.33 |
| Membrane time constant ($\tau$) | Pre-quiet vs. post-quiet | 21.63 | 24.09 | -2.460 | 0.4818 | -9.387 to 4.468 |
|  | Pre-quiet vs. active | 21.63 | 25.98 | -4.352 | 0.2054 | -11.14 to 2.432 |
|  | Post-quiet vs. active | 24.09 | 25.98 | -1.893 | 0.6503 | -10.17 to 6.385 |
| Nº of action potentials<br>(250 pA) | Pre-quiet vs. post-quiet | 41 | 71 | 30 | **0.0076** | -52 to -8 |
|  | Pre-quiet vs. active | 41 | 62 | 21 | **0.0389** | -41 to -1 |
|  | Post-quiet vs. active | 62 | 71 | 9 | 0.4800 | -17 to 34 |
| R input instant ($M\Omega$) | Pre-quiet vs. post-quiet | 205.3 | 139.8 | 65.51 | **0.0079** | 17.67 to 113.3 |
|  | Pre-quiet vs. active | 205.3 | 143.6 | 61.75 | **0.0104** | 14.90 to 108.6 |
|  | Post-quiet vs. active | 139.8 | 143.6 | -3.757 | 0.8962 | -60.92 to 53.40 |
| R input steady state ($M\Omega$) | Pre-quiet vs. post-quiet | 94.70 | 94.46 | -0.2343 | 0.9855 | -25.87 to 25.40 |
|  | Pre-quiet vs. active | 94.70 | 104.1 | 9.394 | 0.4586 | -15.71 to 34.50 |
|  | Post-quiet vs. active | 94.46 | 104.1 | 9.628 | 0.5333 | -21.00 to 40.26 |
| Resting membrane potential (mV) | Pre-quiet vs. post-quiet | -67.24 | -66.71 | -0.534 | 0.8112 | -5.072 to 4.005 |
|  | Pre-quiet vs. active | -67.24 | -62.01 | -5.235 | **0.0319** | -9.980 to -0.4894 |
|  | Post-quiet vs. active | -66.71 | -62.01 | -4.701 | 0.0834 | -10.07 to 0.6643 |
| Rheobase (pA) | Pre-quiet vs. post-quiet | 117.0 | 67.65 | 49.37 | **0.0063** | 14.34 to 84.41 |
|  | Pre-quiet vs. active | 117.0 | 55.56 | 61.47 | **0.0006** | 27.15 to 95.78 |
|  | Post-quiet vs. active | 67.65 | 55.56 | 12.09 | 0.5670 | -29.77 to 53.96 |
| Sag ($\Delta$ mV) | Pre-quiet vs. post-quiet | 18.59 | 7.546 | 11.04 | **0.0002** | 5.422 to 16.66 |
|  | Pre-quiet vs. active | 18.59 | 7.706 | 10.88 | **0.0002** | 5.379 to 16.39 |
|  | Post-quiet vs. active | 7.546 | 7.706 | -0.1591 | 0.9625 | -6.875 to 6.557 |

**Figure 2.**
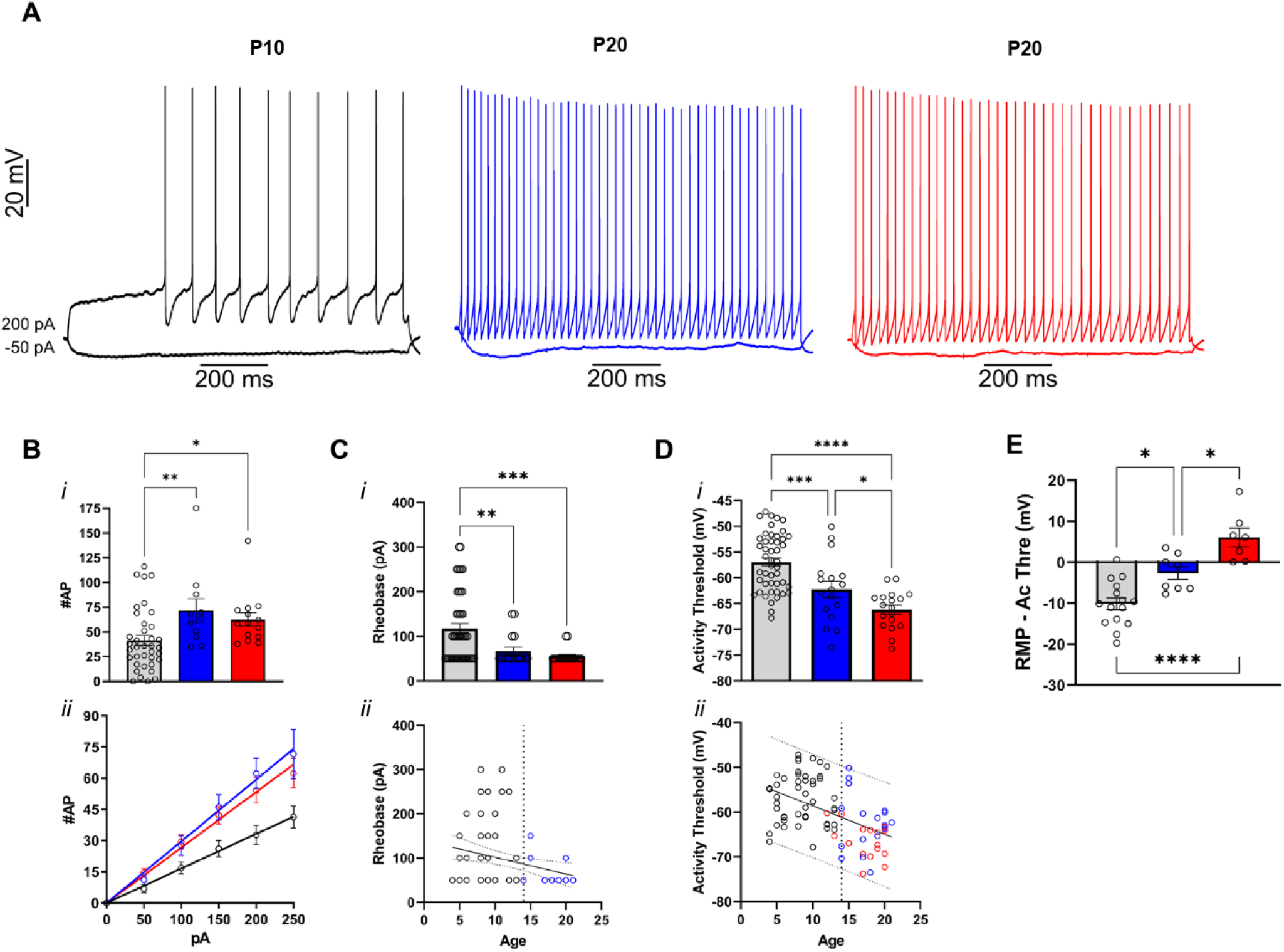
Excitability of fusiform neurons across ages. Gray or black represents pre-hearing quiet neurons, blue post-quiet neurons, and red active neurons. **A**. Representative traces of a quiet neuron from a P10 mouse, a quiet neuron (blue) from a P20 mouse and an active neuron (red) from a mouse of the same age. **Bi**. Number of action potential fired at 250 pA. **Bii**. Number of action potentials fired in response to current injection. **Ci**. Rheobase in pre- and post-hearing neurons and correlation with age (**Cii**). **Di**. The activity threshold and the correlation across age (**Dii**). **E**. Difference between RMP and act. threshold. p <0.05 *, p<0.01 **, p<0.001 ***, p<0.0001 ****.

Pre-hearing quiet neurons also need more current to start firing action potentials, as seen by their bigger rheobases (pre-quiet 117 ± 22.94 pA; post-quiet 67.65 ± 18.05 pA; active 55.56 ± 8.05 pA; Figure 3Ci, Table 1). The bigger rheobase could reflect differences in the voltage where the fusiform neuron starts firing spontaneously, the activity threshold (Leao *et al*., 2012). In fact, pre-hearing quiet neurons had bigger activity thresholds (−56.96 ± 1.63 mV) than post-hearing quiet (−62.25 ± 3.29 mV) and active neurons (−66.20 ± 1.86 mV). In addition, post-hearing quiet neurons also presented a more depolarized activity threshold (one-way ANOVA: p<0.0001; pre-vs. post-quiet p=0.0009; pre-quiet vs. active p<0.0001; post-quiet vs. active p=0.03; Figure 2Di, Table 1). We also found a negative correlation between the activity threshold and postnatal days (Pearson r = -60.66, Table 2; Figure 2Dii). The difference of the RMP and activity threshold was more negative in pre-hearing neurons (mean diff -10.13 ± 3.08, p<0.0001, n=15) than in quiet hearing neurons (mean diff -2.62 ± 3.27, p=0.14, n = 8) and active neurons (mean diff 6.08 ± 5.5, p=0.03, n=7) reflecting the more hyperpolarized RMP and more depolarized activity threshold of pre-hearing quiet fusiform neurons (Figure 2E).

**Table 2.** correlation data statistics. The n =47 of pre-quiet, 17 of post-quiet, and 18 of active. Bold values highlight Person r between 0.5 and 1 or -0.5 and -1.

|  | Intrinsic states | Pearson r | R<sup>2</sup> | p value | 95% confidence interval |
| --- | --- | --- | --- | --- | --- |
| Activity threshold (mV) | Pre-quiet | 0.1375 | 0.0189 | 0.7242 | -0.5795 to 0.7346 |
|  | Post-quiet | -0.1606 | 0.0258 | 0.7309 | -0.8151 to 0.6740 |
|  | Active | **-0.6687** | 0.4471 | **0.0698** | -0.9335 to 0.06805 |
|  | All data | **-0.6063** | 0.3676 | **0.0128** | -0.8473 to -0.1581 |
| AP Amplitude ($\Delta$ mV) | Pre-quiet | **0.7911** | 0.6258 | **0.0111** | 0.2674 to 0.9540 |
|  | Post-quiet | -0.1092 | 0.0119 | 0.8157 | -0.7967 to 0.7016 |
|  | Active | 0.3940 | 0.1552 | 0.3342 | -0.4301 to 0.8599 |
|  | All data | **0.8761** | 0.7676 | **<0.0001** | 0.6725 to 0.9565 |
| AP Latency (s) | Pre-quiet | 0.2348 | 0.0551 | 0.5430 | -0.5086 to 0.7777 |
|  | Post-quiet | 0.4414 | 0.1948 | 0.3215 | -0.4669 to 0.8965 |
|  | Active | 0.4061 | 0.1649 | 0.3182 | -0.4183 to 0.8636 |
|  | All data | -0.3137 | 0.0984 | 0.2368 | -0.7005 to 0.2156 |
| AP Threshold (mV) | Pre-quiet | -0.2515 | 0.0632 | 0.5139 | -0.7846 to 0.4954 |
|  | Post-quiet | -0.0919 | 0.0084 | 0.8446 | -0.7903 to 0.7103 |
|  | Active | 0.0467 | 0.0022 | 0.9126 | -0.6804 to 0.7274 |
|  | All data | -0.4223 | 0.1783 | 0.1032 | -0.7591 to 0.09283 |
| fAHP ($\Delta$ mV) | Pre-quiet | **-0.5289** | 0.2798 | **0.0716** | -0.8829 to 0.2084 |
|  | Post-quiet | 0.3622 | 0.1312 | 0.2123 | -0.5375 to 0.8763 |
|  | Active | **0.5595** | 0.3131 | **0.0746** | -0.2396 to 0.9067 |
|  | All data | **0.5205** | 0.2710 | **0.0194** | 0.03348 to 0.8078 |
| Halfwidth (ms) | Pre-quiet | **-0.8969** | 0.8045 | **0.0010** | -0.9783 to -0.5758 |
|  | Post-quiet | **-0.5098** | 0.2599 | **0.2425** | -0.9125 to 0.3948 |
|  | Active | -0.3377 | 0.1140 | 0.4134 | -0.8420 to 0.4816 |
|  | All data | **-0.8695** | 0.7561 | **<0.0001** | -0.9540 to -0.6571 |
| Max. Rate of rise (V.s<sup>-1</sup>) | Pre-quiet | **0.8859** | 0.7848 | **0.0015** | 0.5387 to 0.9759 |
|  | Post-quiet | 0.1997 | 0.0399 | 0.6677 | -0.6513 to 0.8282 |
|  | Active | **0.5185** | 0.2688 | 0.1880 | -0.2934 to 0.8958 |
|  | All data | **0.9241** | 0.8539 | **<0.0001** | 0.7905 to 0.9737 |
| Membrane time constant ($\tau$) | Pre-quiet | -0.0532 | 0.0028 | 0.8919 | -0.6928 to 0.6333 |
|  | Post-quiet | 0.2650 | 0.0702 | 0.5657 | -0.6097 to 0.8487 |
|  | Active | 0.0975 | 0.0095 | 0.8183 | -0.6520 to 0.7506 |
|  | All data | 0.4400 | 0.1936 | 0.0881 | -0.07119 to 0.7682 |
| R input instant (MΩ) | Pre-quiet | **-0.8145** | 0.6635 | **0.0075** | -0.9596 to -0.3277 |
|  | Post-quiet | **-0.6513** | 0.4242 | **0.1130** | -0.9422 to 0.1997 |
|  | Active | -0.2913 | 0.0848 | 0.4840 | -0.8263 to 0.5202 |
|  | All data | **-0.7723** | 0.5964 | **0.0005** | -0.9169 to -0.4481 |
| R input steady state (MΩ) | Pre-quiet | **-0.5331** | 0.2842 | **0.1395** | -0.8842 to 0.2029 |
|  | Post-quiet | **-0.5860** | 0.3434 | **0.1668** | -0.9291 to 0.2990 |
|  | Active | -0.1784 | 0.0318 | 0.6726 | -0.7845 to 0.6020 |
|  | All data | -0.2109 | 0.0445 | 0.4329 | -0.6397 to 0.3180 |
| Rheobase (pA) | Pre-quiet | 0.3893 | 0.1516 | 0.3003 | -0.3706 to 0.8370 |
|  | Post-quiet | -0.4137 | 0.1711 | 0.3563 | -0.8896 to 0.4930 |
|  | Active | -0.0278 | 0.0008 | 0.9478 | -0.7184 to 0.6904 |
|  | All data | -0.3470 | 0.1204 | 0.1879 | -0.7190 to 0.1796 |
| Sag ($\Delta$ mV) | Pre-quiet | **-0.8261** | 0.6825 | **0.0061** | -0.9623 to -0.3590 |
|  | Post-quiet | -0.2521 | 0.0636 | 0.5854 | -0.8448 to 0.6183 |
|  | Active | -0.3042 | 0.0925 | 0.4638 | -0.8308 to 0.5097 |
|  | All data | **-0.8666** | 0.7509 | **<0.0001** | -0.9529 to -0.6501 |

**Figure 3.**
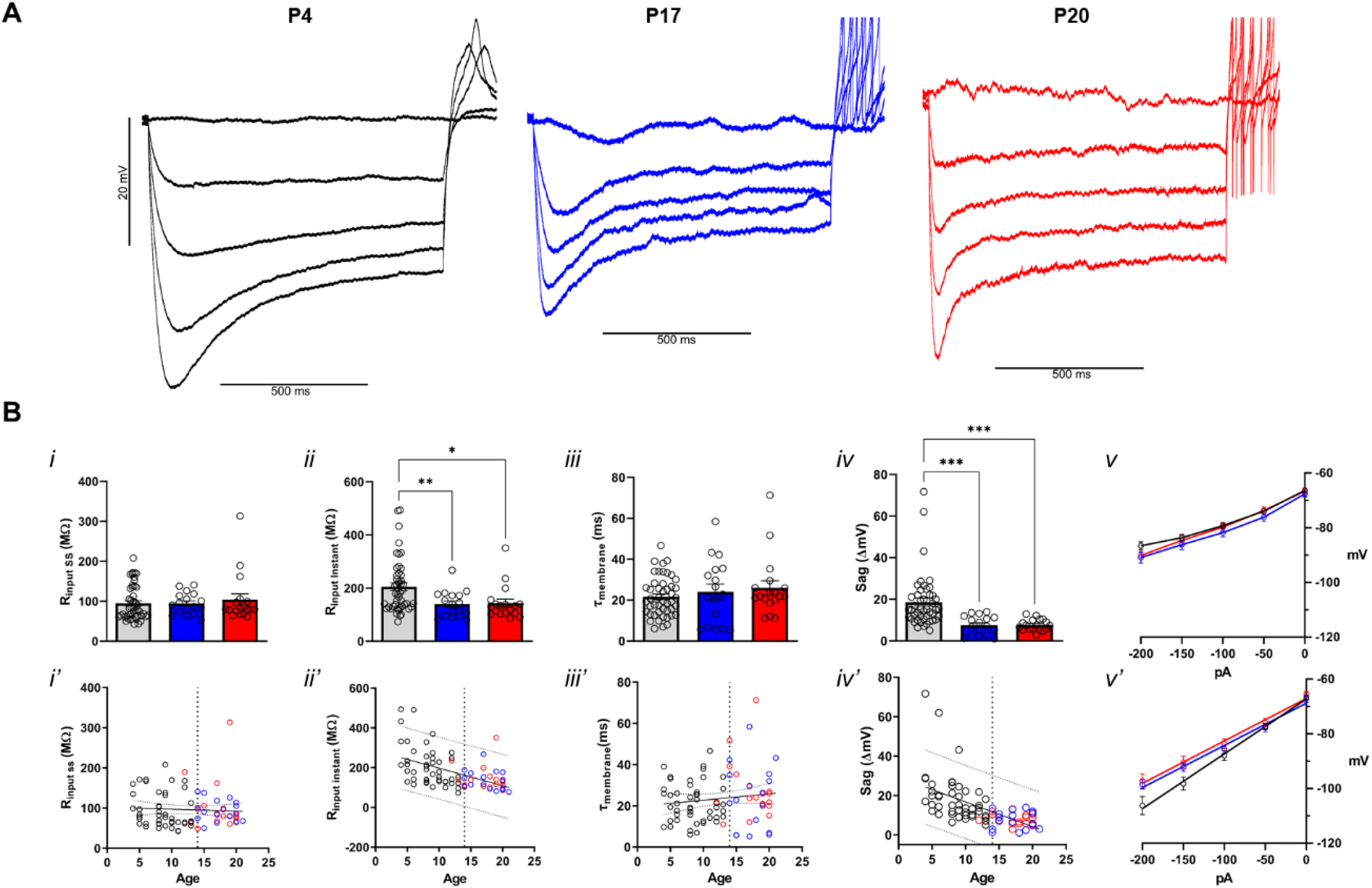
The membrane passive properties of fusiform neurons across ages. Gray or black represents pre-hearing quiet neurons, blue post-quiet neurons, and red active neurons. **A**. Representative traces of fusiform neuron responses to hyperpolarized stimulus from P4 (black), P17 (blue) and P20 (red) mice. **B - i**. Steady state and instantaneous input resistance (**ii**) correlation with age in **ii’. iii**. The time constant of membrane and correlation with ages (**iii’**). **iv**. Depolarization sag and the correlation with age (**iv’**). **v**. VI curve measured at the steady-state and instantly (**v’**). p <0.05 *, p<0.01 **, p<0.001 ***, p<0.0001 ****.

### Subthreshold membrane properties of fusiform neurons change during development

Subthreshold properties of neurons affect their response at rest to synaptic inputs and intrinsic conductances that lead to action potential firing. We compared hyperpolarizing VI relationships of pre-hearing and post-hearing fusiform neurons. Measured at the steady-state, the VI relationships were similar in pre-hearing and post-hearing neurons, showing a small rectification below -80 mV (Figure 3Bv). Membrane input resistance was similar in all groups (pre-quiet 94.70 ± 12.79 MΩ; post-quiet 94.46 ± 13.6 MΩ; active 104.1 ± 30,8 MΩ; one-way ANOVA: p=0.49; pre-vs. post-quiet vs. active: p=0.98 and 0.46; post-quiet vs, active: p=0.53; Figure 3Bi, Table 1) and no significant correlation was observed with age (Pearson r= -0.21,Figure 3Bi’, Table 2). However, the VI relationships measured at the peak hyperpolarization showed that pre-hearing neurons did no present a rectification of the response at more hyperpolarizing potentials (Figure 3Bv’) and a bigger immediate membrane input resistance (pre-quiet 205.3 ± 20.6 MΩ; post-quiet 139.8 ± 23.7 MΩ; active 143.6 ± 30.76; one-way ANOVA: p=0.003; pre-vs. post-quiet vs. active: p=0.008 and 0.01; post-quiet vs. active: p=0.89, Figure 3Bii, Table 1), and a negative correlation with age (Pearson r = - 0.77, Figure 3Bii’, Table 2). Interestingly, the membrane time constant measured near -70 mV did not show a difference between the groups (pre-quiet 21.63 ± 2.75 ms; post-quiet 24.09 ± 7.5 ms; active 25.98 ± 6.6, Figure 3Biii, Table 1), possibly because at this potential, the membrane resistance of all groups is similar as can be seen in Figure 3Bii.

The differences between the input resistance measured at the peak and steady-state suggest that the pre-hearing neurons have a more prominent depolarization sag of the membrane produced by the activation of HCN channels (Ceballos *et al*., 2016). In fact pre-hearing neurons present a more robust depolarization sag than post-hearing neurons (pre-quiet 18.59 ± 3.73 mV; post-quiet 7.55 ± 2.28 mV; active 7.7 ± 1.34 mV; one-way ANOVA: p<0.0001; pre-vs. post-quiet vs. active: p=0.0002; post-quiet vs. active: p=0.96; Figure 3Biv, Table 1) and the sag correlates negatively with age (Pearson r = -0.86; Figure 3Biv’, Table 2). We conclude that changes in the subthreshold membrane conductances during development can contribute to the establishment of quiet and active mature neurons after hearing.

### Action potentials turn progressively faster and bigger during development

We then compared the action potentials of fusiform neurons during the pre- and post-hearing phases. Figure 4Ai show a comparison of representative action potentials from pre-hearing quiet, post-hearing quiet and active neurons, where we can observe that action potentials from pre-hearing neurons are smaller and slower. Accordingly, the amplitude was smaller in pre-hearing neurons (pre-quiet 76.47 ± 4.02 mV; post-quiet 90.63 ± 5.13 mV,; active 96.96 ± 5.19 mV; one-way ANOVA: p<0.0001; pre-vs. post-quiet: p=0.0001; pre-quiet vs. active: p<0.0001; post-quiet vs. active: p=0.135; Figure 4Aii, Table 1), the half-widths were longer (pre-quiet 0,42 ± 0.067 ms; post quiet 0.21 ± 0.06 ms; active 0.19 ± 0.02 ms; one-way ANOVA: p<0.0001; pre-vs. post-quiet p=0.0002; pre-quiet vs. active p<0.0001; post-quiet vs. active p=0.72; Figure 4Aiii, Tabe 1) and the maximum ROR was slower (pre-quiet 570.3 ± 75 V.s^-1^; post-quiet 952.8 ± 90.9 V.s^-1^; active 1036 ± 129.1 V.s^-1^; one-way ANOVA: p<0.0001; pre-vs. post-quiet, and vs. active p<0.0001; post-quiet vs. active p=0.31; Figure 4Biii, Table 1). Both amplitude and maximum ROR also correlated positively with post-natal days (Pearson r = 0.88, and 0.92, respectively, Table 2).

**Figure 4.**
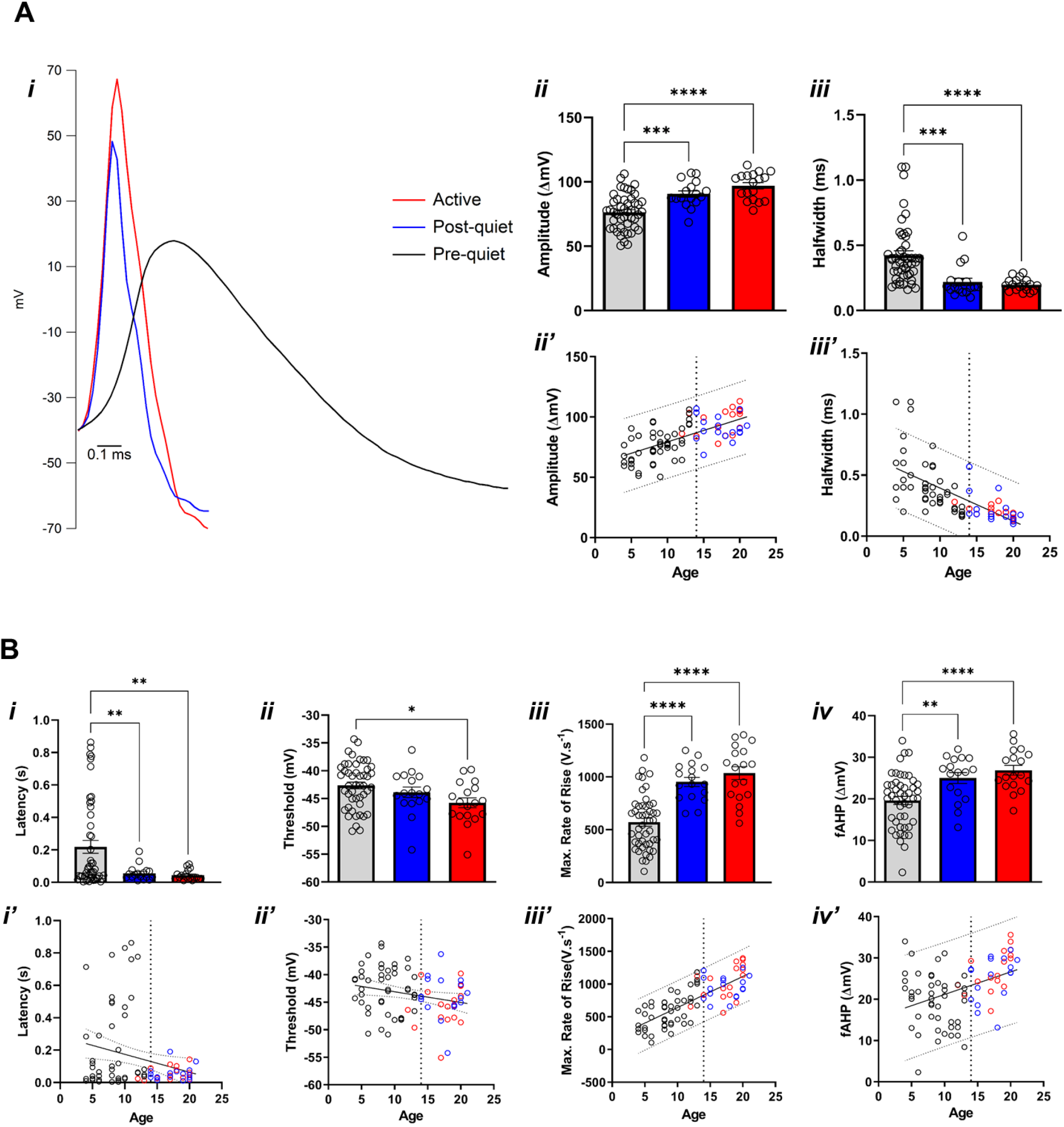
Action potential parameters across ages. Gray or black represents pre-hearing quiet neurons, blue post-quiet neurons, and red active neurons. **A - i**. representative waveforms of action potentials. **ii**. action potential amplitude and the correlation with (**ii’**). **iii**. Half width and correlation with age (**iii’**). **B - i**. Action potential latency and the correlation with age (**i’**). **ii**. Action potential threshold and the correlation with age (**ii’**). **iii**. Maximum rate of rise and correlation with age (**iii’**). **iv**. fAHP and correlation with age (**iv’**). p <0.05 *, p<0.01 **, p<0.001 ***, p<0.0001 ****.

The latency of the first acion potential is longer in pre-hearing quiet neurons (pre-quiet 0.218 ± 0.08 s; post-quiet 0.055 ± 0.02 s; active 0.043 ± 0.01 s; one-way ANOVA: p=0.002; pre-vs post-quiet p=0.007; pre-quiet vs. active p=0.003; post-queit vs. active p=0.86; Figure 4Bi, Table 1). Additionally, action potential thresholds are slightly more hyperpolarized in active neurons (pre-quiet -42.65 ± 1.22 mV; post-quiet -43.88 ± 1.99 mV; active -45.76 ± 1.86 mV; one-way ANOVA: p=0.02; pre-vs. post-quiet p=0.28; pre-quiet vs. active p=0.006; post-quiet vs. active p=0.17; Figure 4Bii, Table 1). We also found a smaller fAHP in quiet pre-hearing neurons (pre-quiet 19.62 ± 1.93 mV; post-quiet 25.00 ± 2.76 mV; active 26.88 ± 2.44 mV; one-way ANOVA: p<0.0001; pre-vs. post-quiet p=0.002; pre-quiet vs. active p<0.0001; post-quiet vs. active p=0.36, Figure 4Biv, Table 1) and a positive correlation of this parameter with age (Pearson r =0.52, Table 2). We conclude that there is a refinement of the action potentials during development to the post-hearing period.

### Principal component analysis (PCA) shows distinct patterns of parameter variations before and after hearing

We performed a PCA of the electrophysiological parameters of the fusiform neurons across the development. We found that most of the variation (∼72%) is contained in the first three components (Figure 5A), with 95% explained in 8 components. Looking at the variation of the different parameters in all components, we can see that most parameters present variations in the first two components, while some others like membrane time constant, action potential threshold, and fAHP in the other four components (Figure 5B)

**Figure 5.**
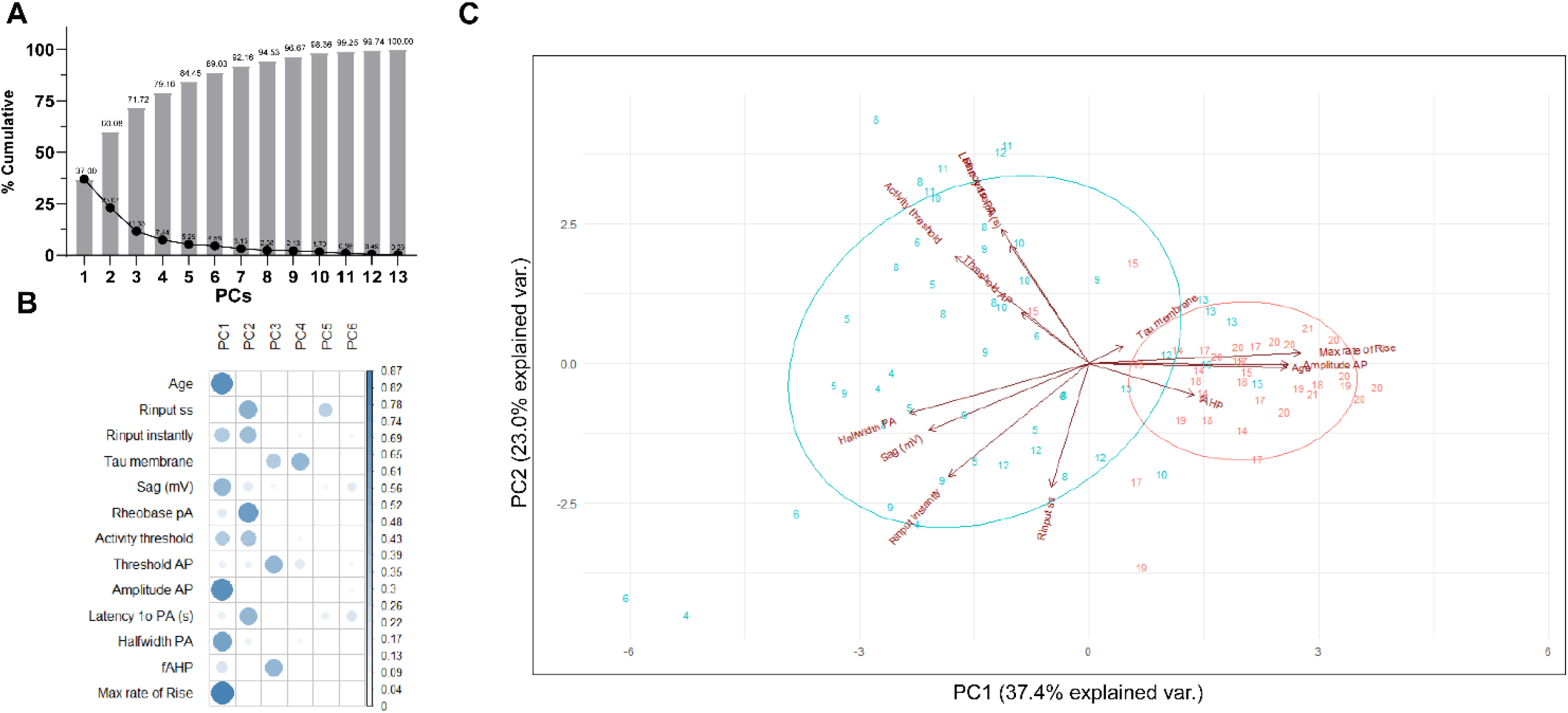
Principal component analysis (PCA) from all data. **A**. The cumulative (bars) and individual (black line) percentage of each principal component (PC). **B**. Contribution of the variables with different PC. **C**. Graph of PC1 vs PC2. The red ellipse represents post-hearing neurons and blue the pre-hearing neurons. Labelled numbers represents animal’s age.

PCA of the first two components shows that most parameter variation occurs during the pre-hearing phase (Figure 5C). The subthreshold parameters Rinput (instantly and on the steady-state) and depolarization sag varied in parallel and opposite the near-threshold parameters AP threshold, activity threshold, latency, and rheobase, in the pre-hearing phase. In the post-hearing phase, we observed an almost parallel correlation of the variation of the AP-related parameters amplitude and ROR.

Comparing the PCAs of pre-earing and post-hearing only, we can identify that several parameters change in parallell in the pre-hearing phase and not after hearing (Figure 6, top). Notably, while immediate and steady-state input resistance showed strong co-variation in the pre-hearing phase, they did not vary in the same direction in the post-hearing period. Latency and activity thresholds vary in parallel during the pre-hearing phase and in opposite directions in the post-hearing phase. Moreover, fAHP that did not show any significant variation in the two components in the pre-hearing phase presented a substantial variation along ROR in the post-hearing phase. Overall we observed less correlations of the parameters in the post hearing phase than in the pre-hearing period. These and other differences show that these parameters evolve differentially in the pre-hearing and post-hearing fusiform neurons.

**Figure 6.**
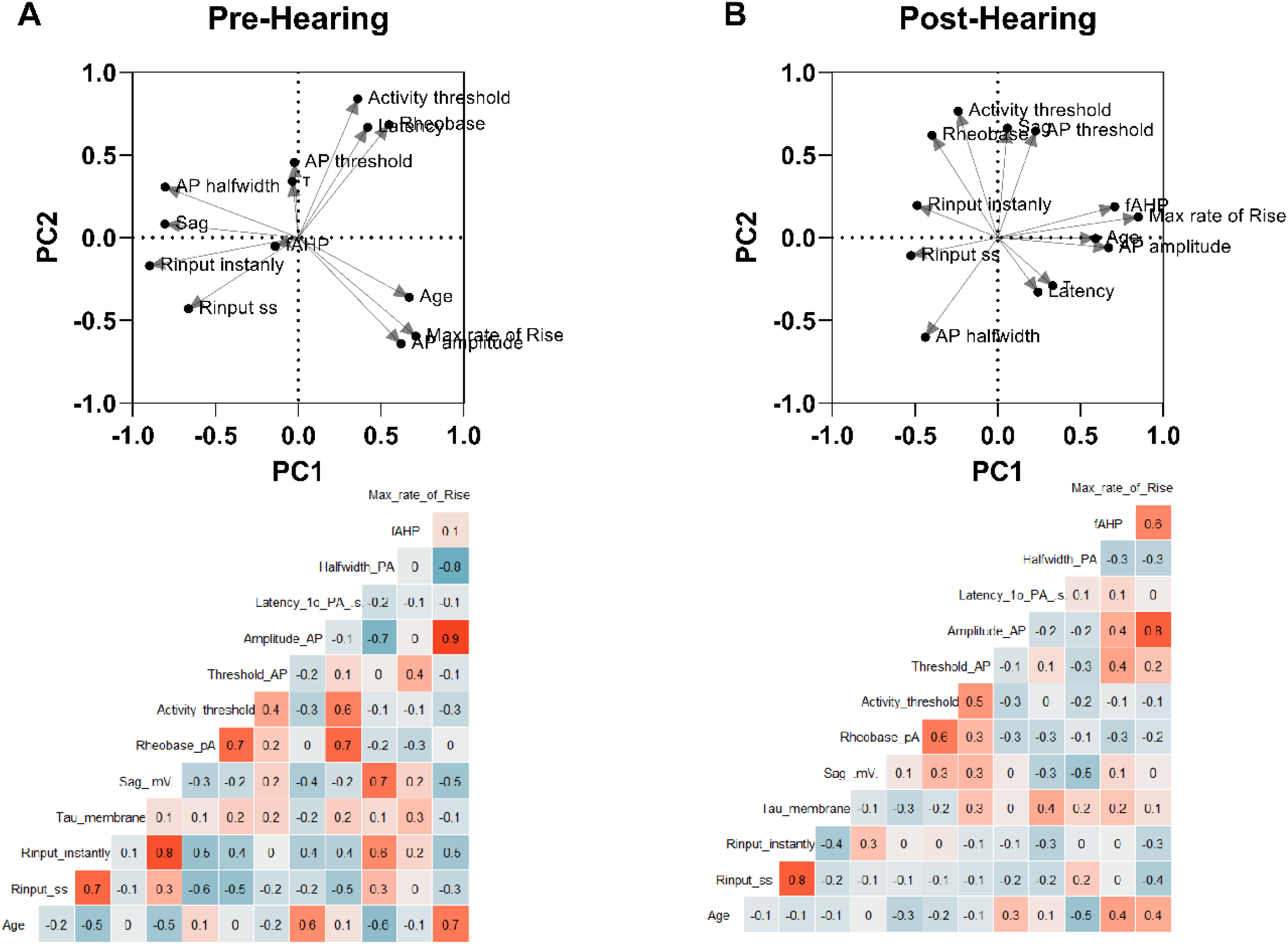
Principal component analysis of pre- and post-hearing neurons. **A**. PC1 vs. PC2 graph from pre-hearing neurons (top) and respective correlogram (bottom). Color intensity represents a degree of correlation. **B**. PC1 vs. PC2 graph from post-hearing neurons (top) and respective correlogram (bottom).

We then analyze the correlations of the variation of the parameters (Figure 6, bottom). We found stronger correlations in the pre-hearing phase than in the post-hearing phase. We found a stronger negative correlation of AP half-width with ROR and amplitude in the pre-hearing phase than in post-hearing phases. However, AP amplitude showed a strong positive correlation between both phases. Activity threshold showed a strong positive correlation with rheobase in both phases and moderate correlation with the threshold. As expected, immediate input resistance showed a strong positive correlation with steady-state input resistance in both phases but a strong negative correlation with the sag and a moderate positive correlation with rheobase and activity threshold only in the pre-hearing phase.

### The persistent sodium current develops during the post-hearing period and creates the active and quiet states

We found that active fusiform neurons appear majorly two days after hearing onset. Our previous works have shown that the active state depends on the sodium persistent current (I_NaP_) expression (Leao *et al*., 2012; Ceballos *et al*., 2016). The I_NaP_ is responsible for creating the activity threshold that initiates action potential firing in potential below the AP threshold (Leao *et al*., 2012). Correlation of activity threshold and postnatal days showed a significant negative correlation with age and a strong hyperpolarization of the activity threshold after P14 (Figure 2D). IV relationships of pre-hearing quiet neurons show that pre-hearing neurons have almost no inward deflection above -60 mV, produced by the development of I_NaP_ (Leao *et al*., 2012; Strazza *et al*., 2021) (Figure 7Bi). The current at -55 mV is significantly bigger in pre-hearing neurons than in post-hearing quiet and active neurons (pre-quiet 106.33 ± 29.29 pA; post-quiet 39.99 ± 46.31 pA; active -85.35 ± 62.25 pA; one-way ANOVA: p<0.0001; pre-vs. post-quiet vs. active: p=0.028 and <0.0001; post-quiet vs. active: p=0.0006; Figura 7Bii, Table 1). Thus, we hypothesized that I_NaP_ increases its expression after hearing, allowing the separation of active and quiet neurons. We then measured I_NaP_ in voltage-clamp as the persistent inward current sensitive to TTX. We found that I_NaP_ is significantly smaller in pre-hearing quiet neurons than in active neurons and post-hearing quiet neurons but there is no differences between post-hearing quiet and active neurons (pre-quiet -26.16 ± 15.28 pA; post-quiet -86.86 ± 29.24 pA; active -161.4 ± 96 pA; one-way ANOVA: p=0.0009; pre-vs. quiet vs. active: p=0.09 and 0.0002; post-quiet vs. active: p=0.06; Figure 7Biv, Table 1). We then conclude that I_NaP_ increases its expression after hearing, creating fusiform neurons’ quiet and active firing modes.

**Figure 7.**
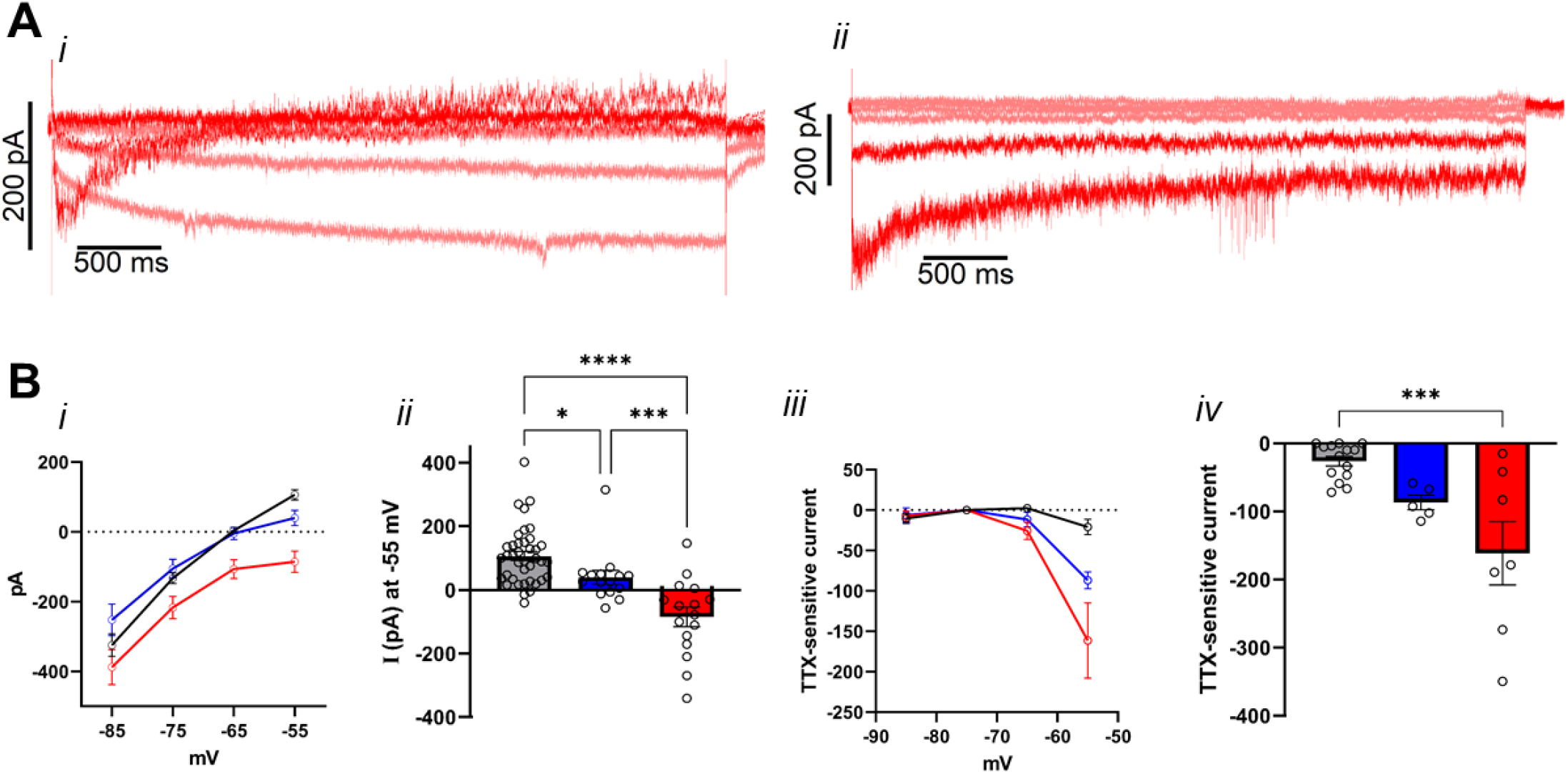
The I_NaP_ in the fusiform neurons during development. Gray or black represents pre-hearing quiet neurons, blue post-quiet neurons, and red active neurons. **A - i**. Representative traces in response to progressive depolarization from -85 mV to -55 mV of an active fusiform neuron. **ii**. TTX-sensitive current from the trace on the left. **B – i**. IV relationships -85 mV to -55 mV. **ii**. Current at -55 mV **iii**. IV relationship of the TTX sensitive current **iv**. TTX-sensitive current at - 55 mV. p <0.05 *, p<0.01 **, p<0.001 ***, p<0.0001 ****.

## Discussion

Several morphological and electrophysiological changes occur during the pre-hearing postnatal development of auditory brainstem neurons to prepare the auditory pathways for the onset of hearing around P12-14. The medial nucleus of the trapezoid body (MNTB) and its specialized terminal from the globular bushy cells from the aVCN, the calyx of Held, has been extensively studied during the pre-hearing to post-hearing periods of development and has been the source of most of the knowledge about the maturation of the brainstem auditory neurons. Most of these studies about the postnatal development of the electrophysiological and synaptic properties of MNTB principal neurons and the calyx of Held showed that the most changes occur before and near hearing onset (Iwasaki & Takahashi, 1998; Taschenberger & Von Gersdorff, 2000; Taschenberger *et al*., 2002; Joshi & Wang, 2002; Leão & Von Gersdorff, 2002; Oleskevich & Walmsley, 2002; Awatramani *et al*., 2005; Youssoufian *et al*., 2005; Leão *et al*., 2005*a*; Nakamura & Takahashi, 2007; Leão & von Gersdorff, 2009; Nakamura & Cramer, 2011). On the other hand, some hearing-dependent changes are observed but, preventing these changes with sound deprivation do not significantly impact the firing and neurotransmission in the MNTB and calyx of Held (Leão *et al*., 2006; Grande *et al*., 2014). Thus, it is accepted that the postnatal development of auditory brainstem neurons is concentrated most in the pre-hearing period and is not significantly affected by the acoustic experience. However, it is known that there is a critical period in humans and rodents around hearing onset that is crucial for full maturation of the auditory system function (De Villers-Sidani *et al*., 2007; Thai-Van *et al*., 2007; Popescu & Polley, 2010; Sun *et al*., 2011; Sharma *et al*., 2016; Zhuang *et al*., 2017; Knipper *et al*., 2020).

Here we studied the development of the electrophysiological properties of the fusiform neurons of the DCN, the primary output neuron of this nucleus, during the pre-hearing and post-hearing periods. Because hyperactivity of fusiform neurons is associated with the development of tinnitus in animal models (Brozoski *et al*., 2002; Kaltenbach *et al*., 2005; Shore *et al*., 2007; Manzoor *et al*., 2013) and tinnitus depends on the establishment of hearing, we believe that changes happening after hearing onset could be relevant for the susceptibility of the DCN to changes in auditory input leading to tinnitus. Tinnitus is a common consequence of deafness, but its development depends on hearing since it is rarely seen in congenitally deaf people (Eggermont & Kral, 2016; Rosing *et al*., 2016). Our more striking finding is that the development of the active state of the fusiform neuron, where it fires action potentials spontaneously (Leao *et al*., 2012), develops after hearing onset. We determined hearing onset at P14 in our animals with ABRs tests and observing the startle to a hand clap and the ear canal opening in pups from litters who knew the exact day of birth. Interestingly, we found that the wave I of the ABR appeared at high thresholds at P13, even though the animal did not react to clapping the hands and had the ear canal sealed. Thus, some acoustic perception could be possible at this age but only to loud sounds, and in a very degraded form, due to the lack of the other downstream waves. The threshold decreased more after P14 with an increased definition of the other waves, suggesting an ongoing maturation of the animal’s hearing. Therefore, the establishment of hearing is not abrupt and improves after opening the ear canal. We found that few active fusiform neurons were detected in the DCN from animals at P12 and P13 (one on each age). The frequency of active neurons increased progressively after hearing onset, peaking 2 to 3 days later. This is strong evidence that the establishment of active fusiform neurons depends on auditory input, and that the active fusiform neuron is important for the computation performed on the DCN and could be relevant for the development of tinnitus.

What is the conductance (or a mix of conductances) in control of the emergence of the post-hearing active neurons? We found that pre-hearing neurons have an activity threshold more depolarized, which can reflect a small expression of the persistent sodium current (I_NaP_), which has been shown to lower the threshold for action potential spontaneous firing and is responsible for the existence of the active state (Leão *et al*., 2012). I_NaP_ is a non-inactivating state of the sodium channels, with a more negative activation than the classic fast inactivating current and is expressed in neurons from many areas of the central nervous system as the hippocampus, thalamus, cerebellum, neocortex, entorhinal cortex, and brainstem regions like cochlear nucleus (Alzheimer *et al*., 1993; Taylor, 1993; Crill, 1996; Ma *et al*., 1997; Ceballos *et al*., 2017*a*, 2017*b*; Hsu *et al*., 2018). Our data suggested that the I_NaP_ expression in the fusiform neuron increased after hearing. Another study found similar results in hippocampal CA1 neurons (Lunko *et al*., 2014). In this study, the I_NaP_ density in neurons from young rats (P12-16) was smaller than in adult rats (P60-75). Also, in mesencephalic trigeminal neurons, there is an increase in I_NaP_ density from P0-6 to P14-17 (Enomoto *et al*., 2018). The expression of the I_NaP_ is related to the expression of auxiliary beta subunits (Qu *et al*., 2001; Aman *et al*., 2009; Bant & Raman, 2010), and the expression of these subunits may be increased in fusiform neurons after hearing. Because the presence of I_NaP_ is essential for lowering the threshold for spontaneous action potential firing (Leão *et al*., 2012), the post-hearing emergence of I_NaP_ is most likely related to the development of the active state of the fusiform neurons.

Besides the emergence of the I_NaP_, post-hearing neurons presented faster, shorter, and bigge action potentials, reflected by their bigger amplitudes, shorter half widths, and faster ROR. Additionally, we observed a significant bigger fast afterhyperpolarization (fAHP) in post-hearing neurons. The variation in the amplitude and ROR were more prominent in the post-hearing neurons, as detected by the PCA. However, the correlation of the parameters with age showed a continuous change during development. Additionally, we found that post-hearing neurons fired more action potentials than pre-hearing neurons when depolarized and had lower action potential thresholds. These changes could be related to changes in voltage-dependent sodium and potassium channels expression. The voltage-gated potassium channel type 3 (K_v3_) is found in different mammalian and avian auditory nuclei areas and mediates the action potential repolarization and modulates fast-spiking neurons. Blockade of K_v3_ channels slows the firing rate of these neurons (Parameshwaran *et al*., 2001; Chambers *et al*., 2017; El-Hassar *et al*., 2019; Choudhury *et al*., 2020). Thus, an increased expression of this class of channels can be related to the shorter action potentials and increased firing of fusiform neurons after hearing.

On the other hand, the bigger amplitude and maximum rate of rise of post-hearing neurons could be related to an increase in the density of voltage-gated sodium channels in the axon initial segment. The fAHP is usually related to calcium-activated potassium channels (K_Ca_) (Louise Faber & Sah, 2003; Adelman *et al*., 2012). An increase in fAHP could be correlated with changes in the expression of these channels or with calcium channel coupling with K_Ca_ channels.

Interestingly, subthreshold parameters present more considerable changes before hearing onset. While immediate input resistance was statistically different in pre-hearing neurons, the membrane resistance measured in the steady-state hyperpolarization was not different, in accordance with the more prominent depolarization sag in the pre-hearing neurons. The sag reflects the activation of the h current by opening the HCN channels in hyperpolarized potentials. These channels have been shown to equalize input resistance in quiet and active fusiform neurons (Ceballos *et al*., 2016) and are probably more prominent in pre-hearing neurons. Interestingly more variation of the immediate input resistance and sag are seen before p10, suggesting that the refinement of these conductances happen earlier in the development. Variations in latency and rheobase are strongly correlated and are more significant in pre-hearing neurons but not in post-hearing neurons. These fluctuations suggest pre-hearing variations of the expression of a low-voltage activated potassium channel that controls latency in fusiform neurons (Kanold & Manis, 1999) but not in post-hearing neurons.

We confirmed that the resting membrane potential of active neurons is more depolarized than pre- or post-hearing quiet neurons. This difference is caused by an increased inwardly rectifying potassium current in quiet neurons (Leao *et al*., 2012). In active neurons, the more depolarized potential crosses the activity threshold set by I_NaP_, generating spontaneous firing (Leao *et al*., 2012; Ceballos *et al*., 2016). in pre-hearing neurons, the low expression of I_NaP_ increases the difference between RMP and activity threshold, making these neurons very difficult to fire, reflecting in their bigger rheobase comparing to post-hearing neurons. Interestingly, the PCA correlations showed that rheobase has a strong positive correlation with activity threshold in both pre-and post-hearing neurons but a strong negative correlation with input resistance only in pre-hearing neurons. Therefore, variations in membrane conductances impact rheobase more substantially in pre-hearing neurons, probably because the changes in the subthreshold conductances are less crucial in post-hearing neurons. We did not investigate the subthreshold currents in this work.

Nonetheless, because of the changes in RMP between active and quiet neurons, we believe that changes in I_Kir_ regulate the firing mode as previously described (Leao *et al*., 2012). Interstingly RMP is similar in pre-hearing and post-hearing quiet neurons, suggesting that their levels of I_Kir_ might be similar and a decrease in I_Kir_ in some neurons after hearing would create the active mode. However, other conductances can affect RMP, like I_h_ and leak potassium and sodium conductances (Leao *et al*., 2012; Ceballos *et al*., 2016), and can be differentially regulated during development. Thus, voltage-clamp experiments characterizing the subthreshold currents during these phases are necessary to answer these questions.

The instant input resistance was bigger in pre-hearing neurons than in post-hearing neurons, but without changes when measured at thw steady-state. Probably slower conductances like the currents mediated by HCN channels (IH) modulate these alterations (Ceballos *et al*., 2016) since we also found a more considerable depolarization sag in pre-hearing neurons. Sag and the instant input resistance are correlated in pre-hearing neurons but not in post-hearing neurons in the PCA, suggesting that, as previously observed, I_h_ equalizes the membrane resistance (Ceballos *et al*., 2016). The PCA also showed a strong negative correlation of action potential half-width with amplitude and maximum rate of rise in pre-hearing neurons but not in post-hearing neurons, suggesting regulated expression of voltage-gated sodium channels and voltage-gated potassium channels in younger animals. Overall our PCA data suggest that several changes in the membrane properties of fusiform neurons occurred before hearing onset, but action potential refinement happened mainly after hearing onset. Additionally more corelations of the parameters were observed in the pre-hearing phase, suggesting that several changes are coordinated in this period, and tend to stabilize after hearing.

## Conclusion

We found that DCN fusiform neurons mature their electrophysiological properties during postnatal development, with changes occurring before and after hearing. Remarkably, the emergence of I_NaP_ and the active state start after hearing onset, suggesting that auditory input could trigger these changes. Because this period is a critical window for proper hearing and tinnitus development, we believe that early hearing loss could impair these changes and impact hearing and tinnitus development. This hypothesis will be tested with hearing deprivation models.

## Acknoledgments

We thank Mr. J. Fernando Aguiar for technical assistance. Research supported by a FAPESP grant (2019/13458-1). NMB is supported by a doctorate scholarship from CNPq. BR receives a undergraduate PBIC scholarship from CNPq. RML holds a CNPq research producitivy scholarship.

